# Partitioning away consciousness: an equal and cross-frequency connectivity analysis from the integration-segregation perspective

**DOI:** 10.64898/2026.06.24.733949

**Authors:** Jose Luis Perez Velazquez, Diego M. Mateos, Richard Wennberg

## Abstract

Derived from previous observations on equal and cross-frequency coupling, we evaluated the proposal that equal and cross-frequency phase synchronization may characterize the integration-segregation perspective of cerebral sensory-motor processing. Using brain recordings obtained in normal conditions and in conditions of diminished sensory input (eyes closed wakefulness, sleep and coma, when there is presumably less functional segregation of sensory-motor processing in neural networks), we assessed potential differences in partitioning of the synchrony state space linked to cross-frequency synchronization. More partitions were found in conditions of decreased sensory input. In addition, there was a less complex synchrony state space in cross-frequency as compared with equal-frequency coupling, in terms of fewer connectivity configurations. These results support the idea that equal-frequency coupling favours integration from multiple brain regions occurring in a complex synchrony state space rich in possible connectivity configurations, whereas cross-frequency coupling contributes to segregation, or localized sensory-motor transformations taking place in specific brain areas. This evidence may contribute to new considerations about the much-discussed role of multi-frequency relations in neuronal activity, and how the structural and functional modular organization of the nervous system is able to generate the coordinated activity needed for consciousness and appropriate cognitive behaviors in complex environments.

## Introduction

The collective coordinated activity of the brain’s cell ensembles results in a variety of neural rhythms spanning all possible frequencies, rhythms whose relations are being vigorously scrutinized using a wide variety of analytical methodologies. To examine those relations, common methods involve the analysis of some sort of synchronization index to assess neural coupling, also termed connectivity, although in reality both of these terms are shorthand for “correlated activity” (see Perez Velazquez and Wennberg (2009) for essentials about the wide field of neuronal coupling/connectivity). Whereas most studies focus on coupling at the same frequency ―1:1 frequency locking― the examination of cross-frequency coupling (*n:m* locking) is attracting much interest as it is thought that these latter relations may play crucial roles in the organization of large-scale networks and functional integration of brain activity (Jirsa and Müller, 2013; Canolty and Knight, 2010).

There have been various specific proposals about the neurophysiological actions of multi-frequency relations in nervous system function, such as the modulation of local synchronization at higher frequencies mediated by lower frequencies (Baptista and Kurths, 2008; Northoff, 2017), or how transient coactivations of localized brain regions may occur at different times during the global propagation of electromagnetic waves throughout the brain (Matsui et al., 2016). A recent study proposed a novel perspective on equal and cross-frequency neural coupling within the framework of integration and segregation of information in cognition (Tononi, 2004), which originated from theoretical considerations regarding the possibility that synchronization is an equivalence relation (Perez Velazquez, 2020). The analysis of phase synchrony in human electrophysiological recordings subsequently resulted in the finding of parcellations of the synchrony state space in the case of cross-frequency phase synchronization (by ‘synchrony state space’ we refer to the space where possible configurations of connections can occur). Hence, the hypothesis associating equal-frequency coupling with integration and cross-frequency coupling with a functional segregation of brain networks was put forward (Mateos and Perez Velazquez, 2025).

To evaluate the validity of this hypothesis, the present study has examined possible differences in the partitioning of the synchrony state space associated with equal and cross-frequency phase synchronization, using human neurophysiological recordings acquired in conditions with different levels of sensory processing, namely in normal awake eyes open states, in awake eyes closed states, in slow wave sleep, and in coma. These distinct levels of the available sensory world engender differences in the need for segregation (and integration too) of the available information: with fewer sensory inputs there should be less segregation. The results obtained provide evidence for the existence of more partitions, or sets, (we will use equivalently the terms partitions or sets throughout the text, with each set of connected signals being one partition) in cross-frequency (*n:m*) connectivity in the case of sensory deprivation. This is associated with a less complex synchrony state space as compared with equal-frequency (1:1) coupling, in that there are fewer possible connectivity configurations in the cross-frequency coupling. The lower complexity associated with more partitions in cross-frequency coupling in turn supports the notion presented by others that the emergence of modularity in biological systems promotes adaptability in organisms by simplifying the search of behaviors in the space of possibilities (Lorenz et al., 2011).

## Materials and Methods

### Neurophysiological recordings

Brain recordings were analyzed from a total of 31 adult subjects. Thirteen were obtained from the open database “EEG Motor Movement/Imagery Dataset” of healthy participants recorded with scalp electroencephalography (EEG) at a sampling frequency of 160 Hz (Schalk et al., 2004). Eleven were 24-hour ambulatory EEG recordings obtained from subjects investigated for unexplained episodic symptoms (e.g., syncope versus seizure; 10 normal, one right temporal lobe epilepsy) at a sampling frequency of 250 Hz. Two were acute and follow-up scalp EEG recordings (sampling frequency 512 Hz) obtained from patients initially in coma, one due to hypoxemic cardiac arrest and the other to sepsis, hypoxia and hypovolemia complicated by left parietal lobe “watershed” infarct; follow-up recording was at 2 weeks and 6 months, respectively (with both patients conscious). The remaining five recordings were described in past studies, all obtained in patients with epilepsy: one magnetoencephalography (MEG) recording and four subjects studied with simultaneous intracranial EEG (iEEG) and scalp EEG (Wennberg, 2010). We used healthy participants wherever possible, but obviously to analyze intracerebral iEEG recordings required the inclusion of epileptic patients. In the latter case, segments with seizures or interictal epileptiform activity were not analyzed. Analogously, segments that contained non-cerebral artefacts like eye blinks were not considered. The number of sensors in each type of recording varied and it is specified, when needed, in the text. In addition, the sampling frequencies varied from 160 to 625 Hz, differences that were taken into consideration for the data analyses.

### Data analyses

A previous study was devoted to the logic and framework for our computation of cross-frequency coupling (Mateos and Perez Velazquez, 2024), therefore we will only highlight very briefly the basic methodology.

Scalp EEG recordings were pre-processed with the current source density (CSD) toolbox (MATLAB implementation (Kayser and Tenke, 2006)) to compute the scalp surface Laplacian or CSD estimates for surface potentials in order to avoid the effects on synchronization measurements related to volume conduction and the common reference electrode used in scalp EEG acquisition. To further diminish signal summation and volume conduction, one half of the sensors of the high-density scalp EEG and MEG recordings were used in analysis, the retained electrodes being equally distributed on the scalp. Variable lengths of the recordings were used for the different analyses, ranging from 1 to 5 minutes, and these segments were divided into consecutive windows ranging from 4 to 30 seconds (details specified in the text as needed).

The analytical procedures start with the computation of the phase synchrony index calculated for all possible pairwise signal combinations, for which we use the standard procedure of estimating phase differences between two signals from the instantaneous phases extracted using the analytic signal concept via the Hilbert transform (Mormann et al., 2000; Pikovsky et al., 2001). To compute the synchrony index, a central frequency of 5 Hz was chosen with a bandpass filter of ±2 Hz. This filter is necessary because to have a clear physical meaning of the instantaneous phase the signal has to be a narrow-band signal (for details on the analytic signal approach see appendix A2 in Pikovsky et al., 2001). The phase synchrony index R between two signals is calculated from the average difference between the phases of the oscillations using a 1-second running window (this window to compute R should not be confused with the window of the signals we use in the rest of the analyses) using the mean phase synchrony statistic, which is a measure of phase locking and is defined as 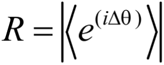 where Δθ is the phase difference between two signals, also termed relative phase: Δθ(t) = θ_1_(t) - θ_2_(t). The more stable the Δθ in the specific time window (1 second), the closer the R value is to 1 (R values are between 0 and 1).

The synchrony indices obtained in this manner for all pairwise combinations of signals in a specific neurophysiological recording gave us a matrix whose entries are the average values of the index R for each pairwise configuration during the time period investigated. From this matrix, a Boolean (binary) connectivity matrix is calculated, with entry of 1 if the corresponding synchrony index is higher than a threshold, and 0 if lower. The threshold is obtained from the mean R given by 20 phase-shuffled surrogates of the original signals. For this purpose, another phase synchrony matrix of the surrogates was obtained by applying the aforementioned method, and each entry in this surrogate matrix was compared with the corresponding entry in the original matrix of the signals, resulting in the binary “connectivity” matrix where a value of 1 is assigned to an entry if the R value of the signals is larger than that of the corresponding surrogates. Please note that we do not use an average of all pairwise surrogate indices to define the threshold, although the results were qualitatively similar if a grand average threshold was used (nonetheless, we think it is more accurate to use the threshold derived from the surrogates for each specific pair of signals).

To compute cross-frequency, or *n:m* phase locking, we follow the same methods described in the previous paragraphs that were specific for 1:1 synchrony. The condition for cross-frequency phase locking is analogous to the equal-frequency synchrony, the difference now is that the relative phase is Δθ*_n,m_*(t) = *n*θ_1_(t) - *m*θ_2_(t), where *n* and *m* are integers, and θ_1,2_ are the phases of the two oscillators (Tass et al., 1998). Our method to compute *n:m* synchrony and the Boolean connectivity matrix then proceeds identically as described above, by extraction of the instantaneous phases of the signals at the required frequency ratios; in our studies we used *n*=1 and *m* ranged from 1 to 4 (the particular *n:m* ratios studied are specified in the Results section as needed). We note that sometimes the ratio 1:4 could not be analyzed because there was no power in the signals at the higher frequency, as power spectra were evaluated prior to the analysis. The only disparity with the 1:1 case is that while the synchrony index matrix is symmetrical for 1:1 it is not so for *n:m* due to the different R values when the phase synchrony is computed in one direction and the reverse; this occurs because the instantaneous phase at frequency *n,* or *m*, is different for signal 1 and signal 2 and thus the phase difference will only be identical if *n=m*. We consider two signals “connected” if they are so in both directions for the reasons detailed in a previous study (Mateos and Perez Velazquez, 2024), consequently in the case of the *n:m* matrix we have to evaluate both entries for each pair of signals in the corresponding binary matrix. Once the binary connectivity matrices were obtained, sets of connected signals were found based on the concept that “if signal 1 is connected to signal 2, and signal 2 is connected to signal 3, then 1 is also connected to 3”. These sets constitute the partitions of the synchrony state space.

To estimate the number of configurations of signal connections we followed identical procedures as detailed in previous studies (Mateos et al., 2017). The statistical significance of the results was evaluated using non-parametric tests, as the data were not normally distributed. Specifically, the Wilcoxon signed-rank test was applied for paired comparisons, and the Kruskal-Wallis test for multiple comparisons, followed by Dunn’s post-hoc test to identify the differing groups.

## Results

The possibility that the number of connectivity configurations differed between equal-frequency (1:1) and cross-frequency (*n:m*) coupling was first investigated. Figure 1 depicts an intuitive explanation of the reason why the number of coupling configurations should be lower as the number of partitions in the synchrony state space increases. As mentioned above, we use the expression ‘synchrony state space’ to refer, in an intuitive manner, to the space where the possible configurations of connections occur.

**Figure 1.**
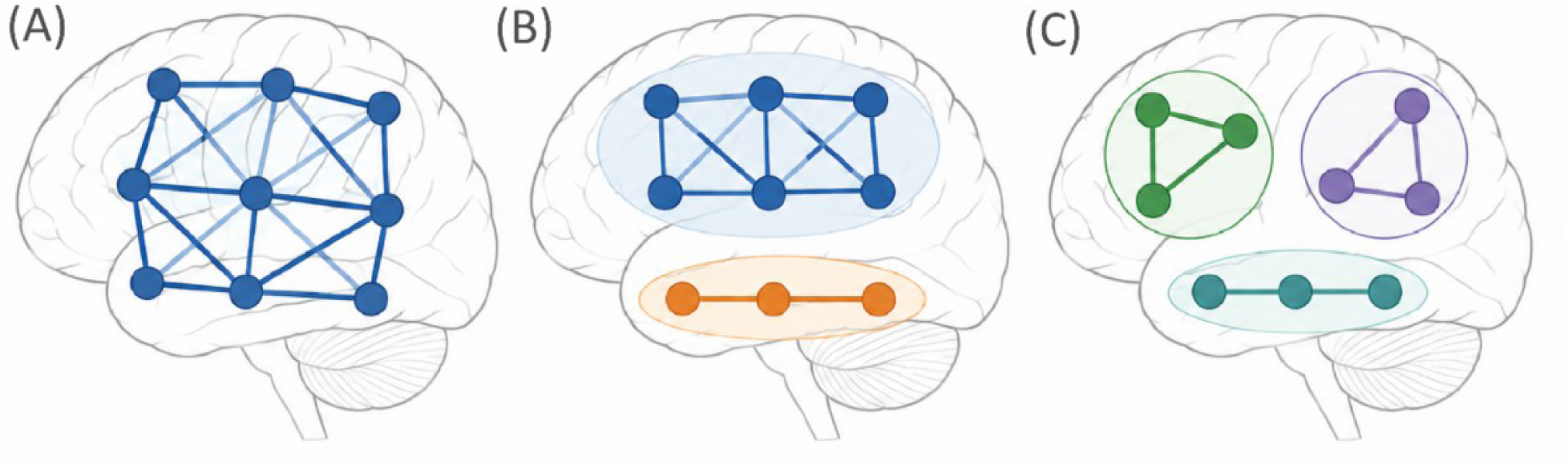
Intuitive demonstration of the relation between lower number of connectivity configurations as the number of partitions of the connectivity state space grows. Assuming coupling only between neighbours, with no partitions (A) there are 18 links, with 2 partitions (B) there are 13 links, and with 3 partitions (C) there are only 8 links. The more links, the more possible configurations of connections.

As anticipated, the number of connectivity configurations was larger in equal-frequency coupling than in cross-frequency coupling in all cases analyzed. Figure 2 illustrates a few specific examples for equal and cross-frequency phase synchrony and the comparison between the total number of configurations in 12 subjects. Similar results were obtained regardless of the state of the subject (awake or asleep) or the type of recording (scalp EEG, iEEG or MEG). The graph of figure 2C includes only those subjects who had the same recording type, because the number of configurations obviously depends on the number of signals, so those with 19-channel scalp EEG were graphed, but the same tendency was observed with any type of recording, as shown in figure 2B for one MEG recording with 144 channels.

**Figure 2.**
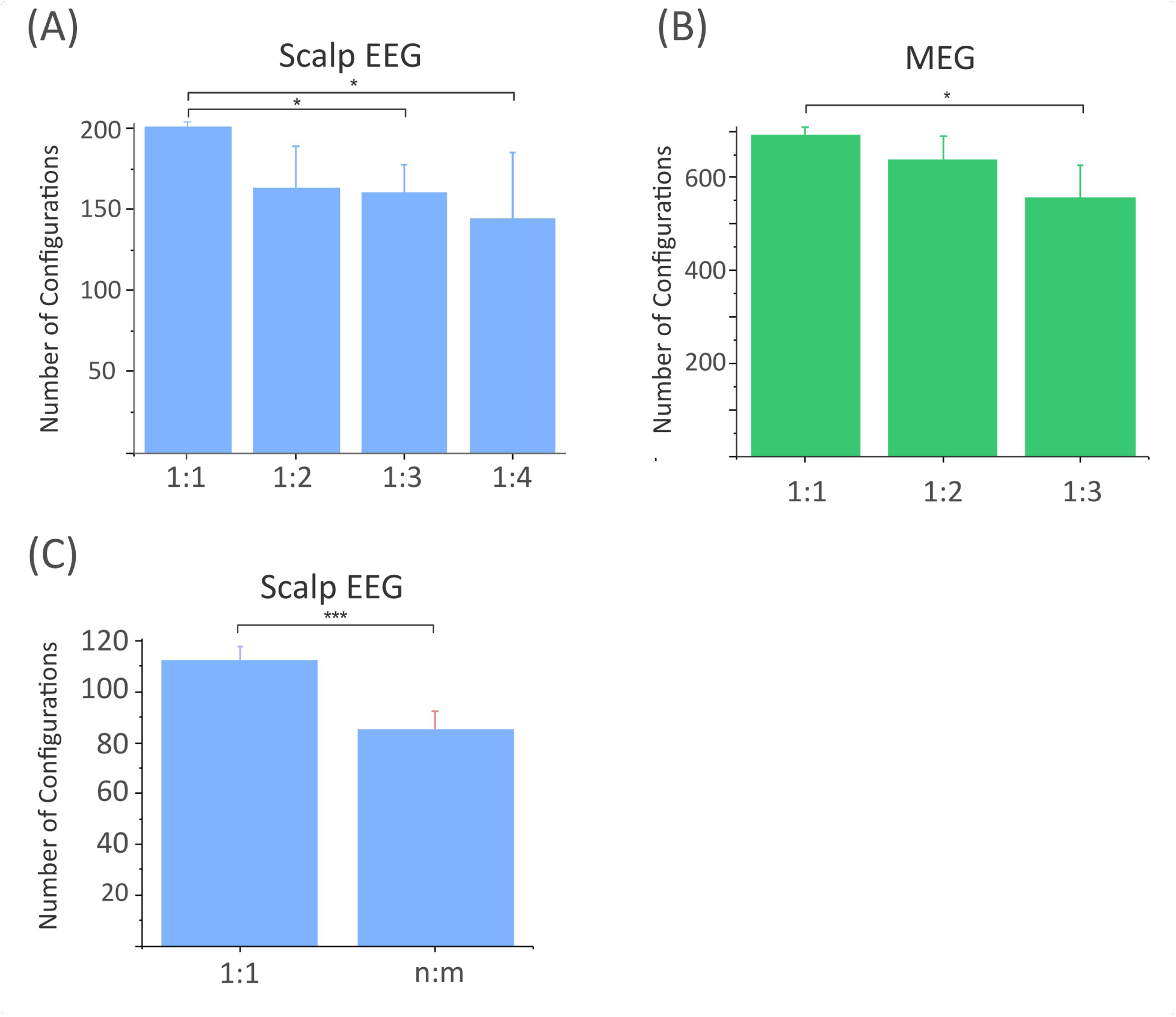
The number of connectivity configurations is lower in cross-frequency coupling. A and B, number of configurations at different locking ratios in one subject recorded with scalp EEG and another with MEG. C, number of configurations for equal and cross-frequency locking (ratios 1:2, 1:3 and 1:4 with central frequency 5 Hz) in 9 subjects recorded with 19-channel scalp EEG. Statistical analysis in A and B used the Kruskal-Wallis test, followed by Dunn’s post-hoc test (*p < 0.05); statistical analysis in C used the Wilcoxon signed-rank test (***p < 0.001)

This result suggests that the state space where the dynamical regimes reside is less complex (and by this we mean it contains fewer possible configurations to explore) in cross-frequency coupling. This makes sense if, as elaborated in the Discussion section, the hypothesis of an association between cross-frequency coupling with the segregation of information, and equal-frequency coupling with the integration of information, holds true. In this case, one would expect a difference in the number of partitions of the synchrony state space between normal cognitive conditions and situations where segregation is diminished. Hence, to examine possible differences, the number of sets (partitions) was evaluated at the three cross-frequency ratios described in the Methods section, in conditions of full wakefulness with eyes open, with eyes closed, in slow wave sleep and in coma. Figure 3 represents the summary of the results comparing wakefulness with sleep. The number of sets was larger during the slow wave sleep state in 13 of the 14 subjects studied, and the results were qualitatively similar whether taking short (4 sec) or long (20 to 30 sec) windows in each recording sample, whose total duration varied from 1 to 5 minutes. Considering all possible comparisons (each subject at three cross-frequency ratios in long and short windows), the number of sets during sleep was greater in 76.2% of the cases. We note that, as previously reported (see figure 4(d) in Mateos and Perez Velazquez, 2024), there is considerable variability in the number of sets in each time window. As an illustration, figure 3A and 3B depict those values in consecutive 4 sec windows of an approximately 1 min recording sample.

**Figure 3.**
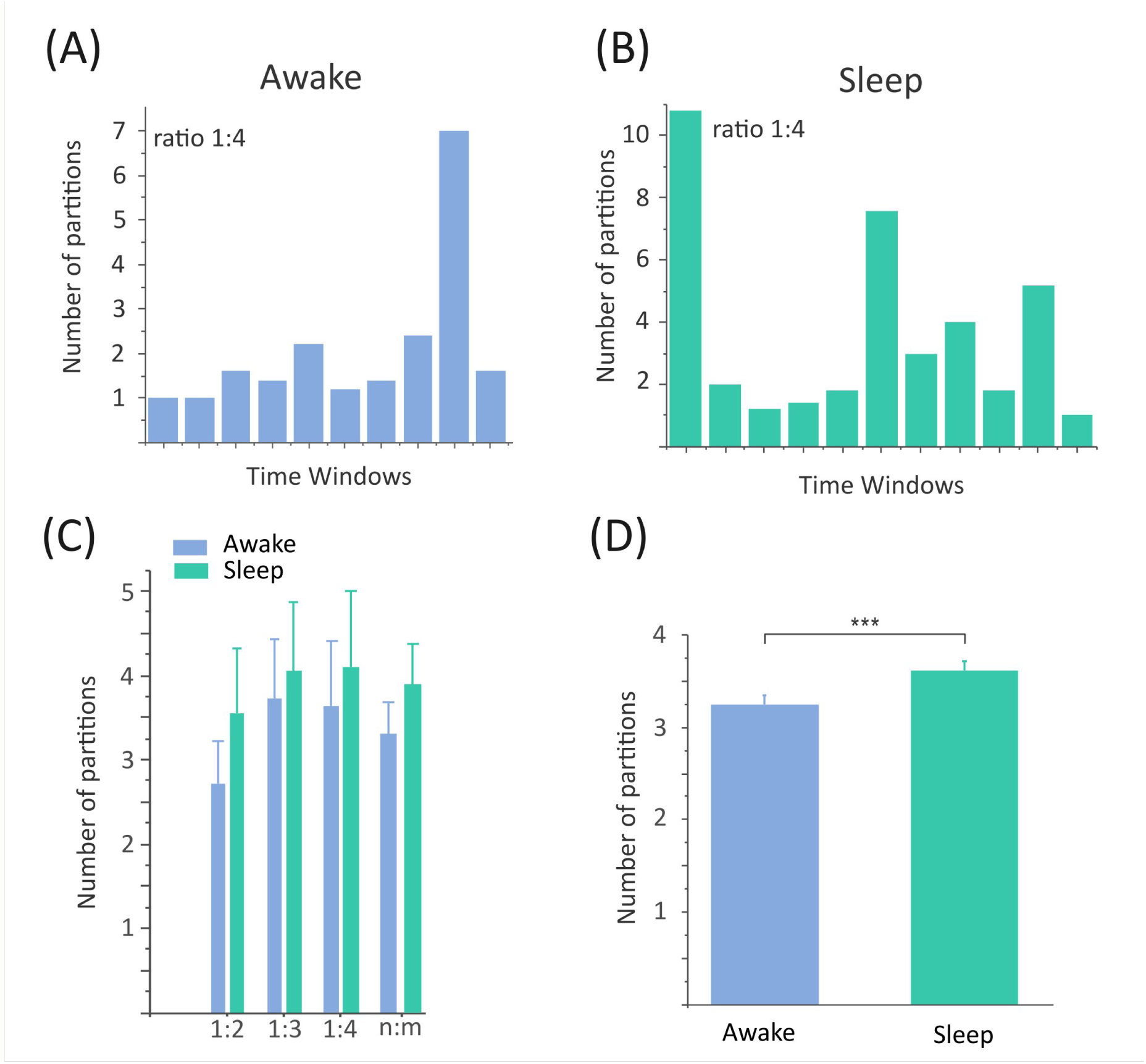
Comparison between the number of partitions in cross-frequency connectivity during awake and slow wave sleep conditions. A and B, each bar represents the number of sets in consecutive windows of 4 seconds in one subject during the awake (A) and slow wave sleep (B) states (ratio 1:4 at central frequency 5 Hz). C, number of sets in one subject during awake and sleep states at the three locking ratios, as well as for the pooled data (n:m bars). D, number of sets during the two states across 14 subjects recorded with 19-channel scalp EEG (***p = 0.0028), Wilcoxon signed-rank test.

**Figure 4.**
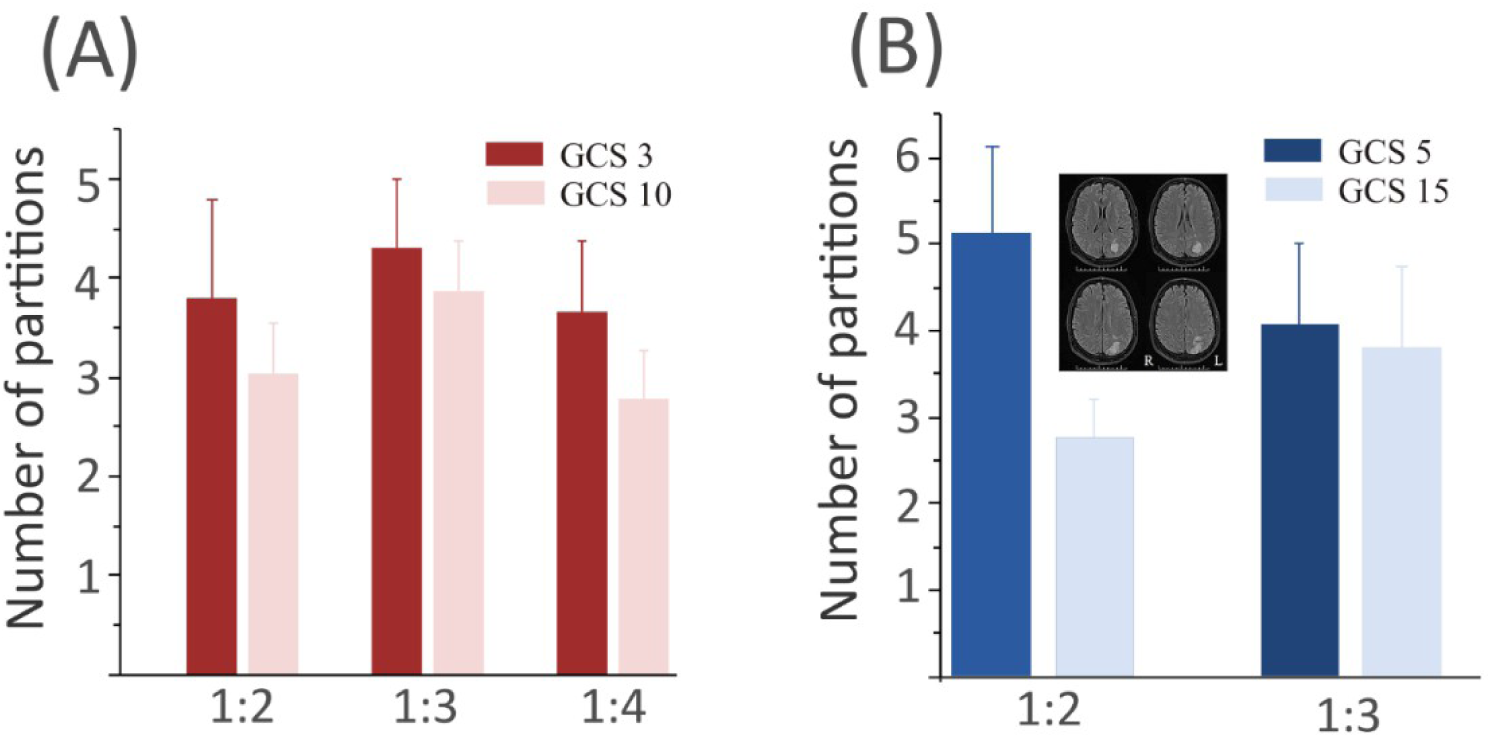
Comparison between the number of partitions in cross-frequency connectivity at the ratios indicated in two patients in coma (dark bars) and after regaining consciousness (light bars). Central frequency = 5 Hz. A, patient 1, post-hypoxemic cardiac arrest. B, patient 2, post-hypoxia/septic shock, complicated by hypovolemic left parietal infarct (brain MRI scan inset). Sagittal and parasagittal parietal channels excluded from analysis in patient 2 due to the extensive structural damage in the region as can be seen in the MRI inset. Glasgow coma scale = GCS (modified GCS-intubated scale used for patient 1, where maximum score = 10; maximum GCS score for non-intubated patient = 15).

The same analysis performed on 14 subjects during the eyes open and eyes closed conditions rendered a larger number of partitions in the latter situation in 8 of the 14 subjects studied, and in 65.1% of the total possible comparisons. There was a tendency for more sets when subjects had their eyes closed, the average being 3.87±0.2 compared with 3.58±0.2 during eyes open wakefulness. Note that these results are not as robust as those comparing wakefulness versus sleep, nevertheless, the same trend emerges: the brain synchrony state space has more partitions during a period with (putatively) less segregation of sensory inputs.

To gain further insight into the relation between the parcellation of the cross-synchrony state space with the level of sensory-motor transformations, the number of partitions in cross-frequency was evaluated in two cases of patients in coma. Although pathological, we used these two cases because the patients recovered from coma and their level of consciousness was longitudinally quantified using the Glasgow Coma Scale (GCS), which offered us a unique opportunity to evaluate the abovementioned relation. Figure 4 displays the results obtained, where it can be seen that the number of sets diminishes with recovery, a result consistent with the previous results indicating that as the need for more segregation of sensory-motor transformations increases, that is, with the return of consciousness, the parcellation of the cross-synchrony state space is reduced.

## Discussion

Following a dynamical feature found in previous studies that indicated a partitioning of the brain synchrony state space into multiple sets of connected cell networks for multi-frequency coupling, we placed those results in the framework of the integration and segregation of sensory-motor processing and evaluated the number of partitions (sets) of brain synchrony state space in normal conditions and in conditions of diminished sensory input. It is reasonable to presume that there would be less functional segregation of sensory-motor processing in the setting of decreased sensory input, whether mild (e.g., removal of visual input) or more complete (e.g., slow wave sleep or coma). More partitions were found in conditions of decreased sensory input, which suggests that there is a relation between multi-frequency connectivity and the need for segregation in sensory-motor processing.

Derived from theoretical considerations regarding the possibility that neural synchronization is an equivalence relation (Perez Velazquez, 2020; Mateos and Perez Velazquez, 2025), a novel perspective on equal and cross-frequency neural coupling was proposed within the framework of integration-segregation of information in cognition (Tononi, 2004). Since there was no parcellation of the synchrony state space in equal-frequency coupling (1:1 synchrony), this was hypothesized to be associated with the integration of information processed from multiple brain regions. On the other hand, the partitioning found in the case of multi-frequency coupling prompted the proposal that it contributes to segregation, or localized processing in specific brain areas. Our present results —that were qualitatively similar regardless of the recording method: scalp EEG, iEEG or MEG— indicating more partitions when the need for sensory-motor transformations is diminished (during eyes closed wakefulness, sleep, or in different stages of coma) supports that proposal. Whereas the equivalence between segregation and clustering of neural nets had been assumed in past studies (Jang et al., 2024), to our knowledge the notion of relating equal and multi-frequency locking with parcellation (clustering) of the neural synchrony state space, and their relation to integration (the former) and segregation (the latter) of neural activity had not been proposed before.

In addition, there was a less complex synchrony state space —in terms of fewer connectivity configurations— in cross-frequency as compared with equal-frequency coupling. Both results are complementary. The “simpler” synchrony state space associated with cross-frequency locking favours the fast processing of computations in localized brain regions, something needed to process a rich variety of sensory-motor transformations coming from the environment. The requirement for fast processing in a complex environment is facilitated by a simpler space where each input/motor command can be processed quickly and efficiently without the need to search through a space with many configurations. As the need for sensory-motor processing decreases (less input or less movement during eyes closed wakefulness, sleep or coma), the (cross-frequency) synchrony space becomes even simpler. It could even be said that the synchronization state space mimics that simplicity of the environment (simplicity due to the decrease in sensory inputs), becoming simpler. In other words, as sensory input is increasingly restricted more and more cross-frequency coupling is reduced, resulting in further fragmentation of the synchrony state space (more partitions); in Figure 1 we can see how as connection links disappear, the number of partitions increases. Moreover, 1:1 coupling, if related to the integration of information as we have proposed, needs a very rich state space with many possible connectivity configurations in order to put together (and make sense of the complexity of) the sensory inputs and motor activity being processed in several brain regions (Perez Velazquez et al., 2019, 2024). Put together, extreme partitioning precludes consciousness.

As to the possible neurophysiology underlying the results, the larger number of partitions present during conditions of decreased sensorium, e.g., after impeding vision or in slow wave sleep or coma, is understandable because the various neural networks within the corresponding sensory cortices that need to be in some sort of coordination —multi-frequency synchronization— while processing the specific inputs arriving at that region, could lose that coordination since there are fewer, or no, inputs to work on, the coordinated regions of cortical activity thus losing their synchrony and thereby becoming independent networks. Hence, when, for instance, four signals recorded from four channels over visual cortex are “working” to process visual inputs (the neural networks underlying those signals are the real “workers”) then it could be the case that the four signals are forming one set in some locking ratio, but upon losing the input they then become independent because there is no need for coordinated activity. It is clear that we cannot determine this scenario from our analyses as more detailed recordings would be needed, particularly intracerebral electrophysiological data recorded from diverse and well-identified neural networks in the different behavioral conditions

As a technical note, whereas we have used throughout the text phase and frequency locking as equivalent, in principle these are not always strictly equivalent. However, in this type of neurophysiological analysis phase locking implies to a large degree frequency locking because the locking condition used, for practical purposes, is that the difference of the phases of the two oscillations be relatively constant within a time window, which translates to the relative constancy of the difference between the frequencies, that is, frequency locking. We focused on synchrony because of the already known fundamental importance of synchronization of neural activity in the functioning of the brain, and particularly differences in the phase of neural oscillations (ter Wal and Tiesinga, 2017). While this is not the place to expound on this, as there are many texts available dealing with the topic, we just note that a multitude of studies have demonstrated the functional significance of phase relations and synchronization in general in nervous system function, perhaps most clearly demonstrated in studies showing the cooperation via synchronization of neural activity in the motor system, whereby the synchronous action potentials arriving to a target neural population ensures maximal impact to spread the activity necessary for specific behaviors (van Wijk et al., 2012).

Our investigation supports the importance of modularity in living systems. In neuroscience specifically, organization in terms of functional and structural modules has been described and discussed profusely. The modular organization of invertebrate and vertebrate nervous systems is well known (Leise, 1990; Galuske et al., 2000) and functional roles for modularity (in terms of neural assemblies), particularly based on re-entrant loops —re-entry of neural activity that could easily cause cross-frequency locking—, have been proposed and investigated, e.g., modules for specific motor programs (Wickens et al., 1994). Modularity is considered to represent an advantageous organization for organisms, providing efficient dynamics in response to fluctuating environments (Sun and Deem, 2007], thus it is not surprising to find it present in the nervous system in order to respond swiftly to variable surroundings. The notion that the emergence of modularity in biological systems promotes adaptability by simplifying the search of behaviors in the space of possibilities (Lorenz et al., 2011) is supported by our result of a simpler synchrony space with fewer connectivity configurations associated with the parcellation during multi-frequency coupling, where the task of searching the whole space of possible neural connectivity configurations is simplified, as there are fewer configurations than during equal-frequency coupling. On another related theoretical aspect, it was shown that systems with large connectivity tend to be unstable whereas there is a tendency towards stability if connectivity is decreased by the arrangement of interactions between the system constituents in “blocks” — using the original term from a paper that dealt with modularity in ecosystems (May, 1972). Consequently, transferring this result in ecosystems to the nervous system, it can be conjectured that neural equal-frequency coupling provides enough instability to the brain, whereas multi-frequency relations stabilize the system.

These more theoretical considerations are important in order to reconcile the need for stability in the brain’s operations with the requirement for a great diversity of activity to process simultaneously many sensory-motor transformations. Therefore, the global pattern of 1:1 locking conceivably provides the large number of connectivity configurations necessary to integrate information processed in many brain regions. This large number of possible connectivity patterns results in the so-called metastability of brain dynamics, which has been discussed for a long time in neuroscience (Rossi et al., 2025), where states are transient due to the wide variety of connectivity configurations emerging from moment to moment to help the organism navigate its surroundings, which we propose to be a fundamental feature of the brain dynamics associated with conscious awareness (Perez Velazquez et al., 2024). On the other hand, multi-frequency locking, associated with a reduced synchrony state space in terms of the number of possible connectivity patterns, may provide the stability needed in the brain areas processing specific sensory-motor transformations. And since the frequencies of neural rhythms evolve over time, the networks can change from one partition (or set) to another, as we found in the analysis of consecutive time windows where the number of sets of coupled brain areas changes continuously, and, accordingly, the composition of those sets change in each time window. This variable composition of parts reflects as well the fluctuations of brain activity that are fundamental for healthy brain function, as decreased variability in neural activity has been related to pathology (Garrett et al., 2013). The dynamic reconfiguration in brain functional connectivity across spatio-temporal scales is thought to be a crucial feature of nervous system dynamics rather than reflecting noise or being a side-effect of ongoing activity (Mengiste and Battaglia, 2026). Overall, these observations further support the idea that the networks of the nervous system are multiscale entities (Betzel and Bassett, 2017) —at the functional and structural levels— which generate variable behaviors through the fluctuations in functional connectivity (Benisty et al., 2024). This behaviour tends to collapse, so to speak, within a small repertoire of possible action-state trajectories in a hypothetical dynamical state space, which may be explained in part by our finding of a less complex state space associated with the segregation of neural activity. The reduced number of possible states to search in order to select one course of action represents an advantage in order to act promptly in the variable environment in which an organism lives. Interestingly, a previous report indicated a Devil’s Staircase dynamical structure in multi-frequency locking during epileptic seizures (Perez Velazquez et al., 2007), and this being a fractal property of complex dynamical systems it can be conjectured that these multiple relations in phase locking are representations of —or are sustaining— the fractal and thus the much-discussed criticality phenomena of neural activity, whereby small perturbations result in chain reactions affecting a large number of neural elements. We are currently addressing these theoretical perspectives from the viewpoint of dynamical system theory.

One limitation of this study is that we evaluated pairwise relations, being the standard in this field. Although some efforts have been devoted to assess the synchronization of multiple units, e.g., triplets (Kralemann et al., 2013), exploring these relatively complex synchrony patterns will require more advanced analytical methods than those we have used. Nevertheless, we were able to determine, based on pairwise correlations, groups of “connected” signals (the quotation marks refer to what was mentioned in the first paragraph of the Introduction regarding measures of synchrony). Another complication arises from the high variability in the number of partitions in each time window, something that a glance at figure 3A and 3B makes clear. The outcome of this variability is that to obtain a statistical significance with p<0.05 is not always possible. What we have documented is a clear trend towards more parcellation —in multi-frequency connectivity of a hypothetical synchrony state space— being associated with diminished sensory-motor processing, when presumably there is less need for the segregation of the information processed by brain circuits. We thus propose that multiple-locking ratios are related to the segregation aspect of brain dynamics, whereas equal-frequency coupling is associated with the integration side. In short, integration via 1:1 coupling requires a complex space with many coupling configurations to solve problems and move around in the world, whereas segregation in diverse cross-frequency relations results in a simpler space where each input/motor command can be processed rapidly and efficiently without a need to search through a space with many connectivity patterns. As the need for sensory-motor transformations decreases (less sensory input or less movement), the cross-frequency locking state space becomes even simpler. Our observations are obviously not a complete story but represent a preliminary perspective that, to our knowledge, had not been advanced before with respect to cross-frequency coupling, opening up avenues to consider how the structural and functional modular organization of the nervous system generates the coordinated activity necessary for appropriate sensory-motor processing.

## Author contributions

JLPV, DMM and RW conceptualized the study and acquired the data. JLPV wrote the original manuscript draft. All authors critically reviewed the manuscript.

## Data availability

The data analyzed during the current study are available from the corresponding authors upon reasonable request.

## References

Baptista, M. S., and Kurths, J. (2008) Transmission of information in active networks. Phys. Rev. E Stat. Nonlin. Soft Matter Phys. 77:026205. doi: 10.1103/PhysRevE.77.026205

Benisty, H., Barson, D., Moberly, A. H., Lohani, S., Tang, L., Coifman, R. R., et al. (2024) Rapid fluctuations in functional connectivity of cortical networks encode spontaneous behavior. Nat. Neurosci. 27, 148–158. doi: 10.1038/s41593-023-01498-y

Betzel, R. F., and Bassett, D.S. (2017) Multi-scale brain networks. Neuroimage 160, 73–83. doi: 10.1016/j.neuroimage.2016.11.006.

Galuske, R. A., Schlote, W., Bratzke, H., and Singer, W. (2000) Interhemispheric asymmetries of the modular structure in human temporal cortex. Science 289, 1946–1949. doi: 10.1126/science.289.5486.1946

Garrett, D. D., Samanez-Larkin, G. R., MacDonald, S. W., Lindenberger, U., McIntosh, A. R., and Grady, C. L. (2013) Moment-to-moment brain signal variability: a next frontier in human brain mapping? Neurosci. Biobehav. Rev. 37, 610–624. doi: 10.1016/j.neubiorev.2013.02.015

Jang, H., Mashour, G. A., Hudetz, A. G., and Huang, Z. (2024) Measuring the dynamic balance of integration and segregation underlying consciousness, anesthesia, and sleep in humans. Nat. Commun. 15:9164. doi: 10.1038/s41467-024-53299-x

Jirsa, V., and Müller, V. (2013) Cross-frequency coupling in real and virtual brain networks. Front. Comput. Neurosci. 7:78. doi: 10.3389/fncom.2013.00078

Kayser, J., and Tenke, C. E. (2006) Principal components analysis of Laplacian waveforms as a generic method for identifying ERP generator patterns: I. Evaluation with auditory oddball tasks. Clin. Neurophysiol. 117, 348–368. doi: 10.1016/j.clinph.2005.08.034

Kralemann, B., Pikovsky, A., and Rosenblum, M. (2013) Detecting triplet locking by triplet synchronization indices. Phys. Rev. E Stat. Nonlin. Soft Matter Phys. 87:052904. doi: 10.1103/PhysRevE.87.052904

Leise, E. M. (1990) Modular construction of nervous systems: a basic principle of design for invertebrates and vertebrates. Brain Res. Rev. 1, 1–23. doi: 10.1016/0165-0173(90)90009-d

Lorenz, D. M., Jeng, A., and Deem, M. W. (2011) The emergence of modularity in biological systems. Phys. Life Rev. 8, 129–160. doi: 10.1016/j.plrev.2011.02.003

Luo, Z., Yin, E., Zeng, L. L., Shen, H., Su, J., Peng, L., et al. (2024) Frequency-specific segregation and integration of human cerebral cortex: An intrinsic functional atlas. iScience 27:109206. doi: 10.1016/j.isci.2024.109206

Mateos, D. M., and Perez Velazquez, J. L. (2025) Perspective on equal and cross-frequency neural coupling: Integration and segregation of the function of brain networks. Phys. Rev. E Stat. Nonlin. Soft Matter Phys. 111:014408. doi: 10.1103/PhysRevE.111.014408

Mateos, D. M., Wennberg, R., Guevara, R., and Perez Velazquez, J. L. (2017) Consciousness as a global property of brain dynamic activity. Phys. Rev. E Stat. Nonlin. Soft Matter Phys. 96:062410. doi: 10.1103/PhysRevE.96.062410

May, R. M. (1972) Will a large complex system be stable? Nature 238, 413–414. doi: 10.1038/238413a0

Matsui, T., Murakami, T., and Ohki, K. (2016) Transient neuronal coactivations embedded in globally propagating waves underlie resting-state functional connectivity. Proc. Natl. Acad. Sci. U. S. A. 113, 6556–6561. doi: 10.1073/pnas.1521299113

Mengiste, S. A., and Battaglia, D. (2026) Dynamic functional connectivity resolves brain integration-segregation trade-off under costly links. arXiv [Preprint]. arXiv:2604.11608

Mormann, F., Lehnertz, K., David, P., and Elger, C. E. (2000) Mean phase coherence as a measure for phase synchronization and its application to the EEG of epilepsy patients. Physica D 144, 358–369. doi: 10.1016/S0167-2789(00)00087-7

Northoff, G. (2017) “Paradox of slow frequencies” – Are slow frequencies in upper cortical layers a neural predisposition of the level/state of consciousness (NPC)? Conscious. Cogn. 54, 20–35. doi: 10.1016/j.concog.2017.03.006

Perez Velazquez, J. L., Garcia Dominguez, L., and Wennberg, R. (2007) Complex phase synchronization in epileptic seizures: evidence for a devil’s staircase. Phys. Rev. E Stat. Nonlin. Soft Matter Phys. 75:011922 doi: 10.1103/PhysRevE.75.011922

Perez Velazquez, J. L., and Wennberg, R. (2009) Coordinated Activity in the Brain: Measurements and Relevance to Brain Function and Behaviour. Berlin: Springer.

Perez Velazquez, J. L., Mateos, D. M., and Guevara Erra, R. (2019) On a simple general principle of brain organization. Front. Neurosci. 13:1106. doi: 10.3389/fnins.2019.01106

Perez Velazquez, J. L. (2020) On the emergence of cognition: from catalytic closure to neuroglial closure. J. Biol. Phys. 46, 95–119. doi: 10.1007/s10867-020-09543-8

Perez Velazquez, J. L., Mateos, D. M., Guevara, R, and Wennberg, R. (2024) Unifying consciousness theories with MaxCon: maximizing configurations of the brain web. OSF [Preprint]. doi:10.31219/osf.io/af4n5

Pikovsky, A., Rosenblum, M., and Kurths, J. (2001) Synchronization: A Universal Concept in Nonlinear Sciences. Cambridge: Cambridge University Press.

Rossi, K. L., Budzinski, R. C., Medeiros, E. S., Boaretto, B. R. R., Muller, L., and Feudel, U. (2025) Dynamical properties and mechanisms of metastability: A perspective in neuroscience. Phys. Rev. E Stat. Nonlin. Soft Matter Phys. 111:021001. doi: 10.1103/PhysRevE.111.021001

Schalk, G., McFarland, D. J., Hinterberger, T., Birbaumer, N., and Wolpaw, J. R. (2004) BCI2000: a general-purpose brain-computer interface (BCI) system. IEEE Trans. Biomed. Eng. 51, 1034–43. doi: 10.1109/TBME.2004.827072

Sun, J., and Deem, M. W. (2007) Spontaneous emergence of modularity in a model of evolving individuals. Phys. Rev. Lett. 99: 228107. doi: 10.1103/PhysRevLett.99.228107

Tass, P., Rosenblum, M. G., Weule, J., Kurths, J., Pikovsky, A., Volkmann, J., et al. (1998) Detection of *n*:*m* phase locking from noisy data: Application to magnetoencephalography. Phys. Rev. Lett. 81:3291. doi: 10.1103/PhysRevLett.81.3291

ter Wal, M., and Tiesinga, P. H. (2017) Phase difference between model cortical areas determines level of information transfer. Front. Comput. Neurosci. 11:6. doi: 10.3389/fncom.2017.00006

Tononi, G. (2004) An information integration theory of consciousness. BMC Neurosci. 5:42. doi: 10.1186/1471-2202-5-42

van Wijk, B. C. M., Beek, P. J., and Daffertshofer, A. (2012) Neural synchrony within the motor system: what have we learned so far? Front. Hum. Neurosci. 6:252. doi: 10.3389/fnhum.2012.00252

Wennberg, R. (2010) Intracranial cortical localization of the human K-complex. Clin. Neurophysiol. 121, 1176–1186. doi: 10.1016/j.clinph.2009.12.039

Wickens, J., Hyland, B., and Anson, G. (1994) Cortical cell assemblies: a possible mechanism for motor programs. J. Mot. Behav. 26, 66–82. doi: 10.1080/00222895

